# Habitat differences filter functional diversity of low dispersive microscopic animals

**DOI:** 10.1101/2020.09.22.308353

**Authors:** Alejandro Martínez, Guillermo García-Gómez, Álvaro García-Herrero, Nuria Sánchez, Fernando Pardos, Andrés Izquierdo-Muñoz, Diego Fontaneto, Stefano Mammola

## Abstract

Whereas the study of patterns of distribution of microscopic animals has long been dominated by the ubiquity paradigm, we are starting to appreciate that microscopic animals are not as widespread as previously thought and that habitat preferences may have a strong role in structuring their patterns of occurrence. However, we still ignore to what extent and through which mechanisms the environment selects for specific communities or traits in microscopic animals. This gap is partly due to the lack of data on the relevant traits of many species, and partly because measuring environmental variables at an appropriate resolution may be problematic.
We here overcome both issues by analysing the functional space of marine mite communities living in a sea-grass (*Posidonia oceanica*) meadow across two habitats: the leaves and the matte. The strictly benthic lifestyle and the conserved morphology of mites allow for unambiguous characterization of their functional traits, while the discrete nature of the two habitats alleviates the uncertainty in their ecological characterization.
Our results show that habitat filters the distribution of certain traits favouring a higher diversity, dispersion, and evenness of functional traits in the matte than in the leaves. We further observed temporal variations in the functional diversity of communities, potentially following the seasonal renovation and decay of seagrass leaves. However, in spite of the stark ecological differences between the two habitats and across seasons, the filtering effect is partial and affects mostly relative species abundances.
We conclude that in other microscopic organisms, habitat filtering might appear even more subtle especially if they are capable of long distance dispersal or occur in ecological systems where environmental variables vary continuously or fluctuate through time. Our study therefore emphasises the need of moving from a merely taxonomical toward a functional view of ecological studies of microscopic organisms if we want to achieve a mechanistic understanding of their habitat and distribution patterns.

**Data availability statement:** Raw data and R script to generate the analyses will be deposited in a public repository upon acceptance.

## Introduction

It is unlikely to see buffaloes grazing on the sea surface or whales gliding in the sky (Adams, 1984). However, as the body size of animals decreases, the probability increases of encountering them in places where *they are not supposed to be*. This is because the realised niche of a microscopic animal—namely, where it can be actually found—can extend well beyond the set of abiotic conditions that allow positive population growth rates (Grinnellian niche). These broad ecological ranges are more frequent amongst microscopic animals possessing traits that facilitate long distance dispersal such as dormancy, long term viability, and parthenogenesis (Fontaneto & Hortal, 2013, Fontaneto, 2019). Similar traits are found, for example, in many species of nematodes (Fonseca & Netto, 2015), rotifers (Fontaneto, Barraclough, Chen, Ricci, & Herniou, 2008), and tardigrades (Bartels, Kaczmarek, Rozkowska, & Nelson, 2020; Kaczmarek, Michalczyk, & McInnes, 2015). In comparison, some lineages of microscopic organisms are specialised to thrive within narrow ranges of environmental conditions like caves (Mammola et al., 2020), mountain summits (Hoschitz & Kaufmann, 2004), hydrothermal vents (Zeppilli et al., 2018), and deep terrestrial subsurface habitats (Borgonie et al., 2011). Many of these animals evolved distinct and often convergent traits for these specific conditions. Quintessential examples are microscopic annelids and copepods specialised to feed in the chemocline of certain aquatic caves (Martínez et al., 2019; Worsaae et al., 2019); or mouthless species of nematodes and flatworms living in strict association to prokaryotic symbiont in anoxic marine sediments (Ott, Rieger, Rieger, & Enderes, 1982).

The corollary of these examples is that not only the body size but also the presence of certain traits and the interaction between them and the environment determines the ecological range of microscopic organisms. This is nothing new, as this idea was already grasped in the original formulation of the “*everything small is everywhere*” paradigm, which included the postil “…*but the environment selects*” (Baas-Becking, 1934; Bass & Boenigk, 2011). So we now stand to a point where we know that even broadly distributed and apparently generalist species may not be actually so widespread and tolerant when their habitat preferences are taken into account (or, in other words, that the density of individuals across the distribution range of a given species is not homogeneous as it varies across habitats). But, unfortunately, this filtering effect has proven difficult to quantify, partly due to the lack of data on the relevant traits of many microscopic animals (Giere, 2008) and partly due to the intrinsic problem of measuring relevant environmental variables at appropriate resolutions (Levin, 1992; Potter, Arthur Woods, & Pincebourde, 2013) overestimating the Grinnellian niche (Soberón & Nakamura, 2009). These two issues have challenged all community-level studies that have so far attempted to directly link functional traits of microscopic animals and their distribution patterns at the relevant scale (Fontaneto et al. 2011). In other words, we ignore to what extent and through which mechanisms the environment selects for specific communities and their traits.

We here set to examine the effect of habitat on the distribution of microscopic animals by comparing the multidimensional functional space (Blonder, Lamanna, Violle, & Enquist, 2014; Blonder et al., 2018) of assemblages of mites dwelling on a seagrass [*Posidonia oceanica* (L.)] meadow in the Mediterranean—a marine plant with a well-studied architecture and growth pattern (Molenaar, Barthélémy, De Reffye, Meinesz, & Mialet, 2000). Due to their strictly benthic life mode and easy-to-measure external traits with a clear functional meaning, marine mites are an excellent model system for a similar analysis. Furthermore, the patchy distribution of seagrass within meadows provides independent replicates of discrete habitats, the leaves versus the matte (i.e., the grid formed by rhizomes, roots, and trapped particles). Because these two habitats present different environmental conditions and availability of food, we expect that they will filter different mites from the pool of species present in the meadow. We expect that this filter will be evidenced in the community traits, favouring the dominance of more specialised phytophagous or epiphytes feeder species in the leaves, and limiting the presence of generalistic detritivorous species to the matte. We therefore hypothesise that i) at the community level, there should be higher diversity, dispersion, and evenness of functional traits in the matte than in the leaves. As a corollary of the previous hypothesis, we also expect that ii) at the species level, the higher diversity of traits in the matte will be reflected by the presence of more functionally original species. Furthermore, the annual phenological changes due to the seasonal renovation and decay of seagrass leaves affects nutrient availability (Drew, 1978; Zupo, Buia, & Mazzella, 1997). So, we also hypothesize iii) temporal variations in the functional diversity of mite communities following the annual cycle of *P. oceanica*, particularly on the leaves.

## Material and Methods

### Model organism

The model organisms selected for this study are marine mites of the family Halacaridae (subsequently referred to as marine mites), a lineage of microscopic arachnids that colonized the ocean from a terrestrial ancestor around 270 million years ago, radiating in different types of marine habitats (Pepato, Vidigal, & Klimov, 2018). Due to this terrestrial origin, the body plan of the group is constrained, being all forms strictly restricted to benthic habitats. The impossibility of marine mites to swim or disperse by any other means than crawling in direct contact with the substrate, ensures that the species found in each sample belong to the local community. This feature places marine mites among those with a realised niche that is smaller than the potential Grinnellian niche, even if they are microscopic: not all available habitats in an area are colonised, and the animals are not found in habitats that cannot sustain viable populations. Furthermore, the presence of a hard, hydrophobic cuticle allows for a precise measurement of morphological traits even in fixed material, reducing measurement errors. Finally, the conserved morphology ensures unequivocal homology assessment of the functional traits. These three properties— dispersal exclusively by crawling, hard cuticle, and conserved morphology—make marine mites ideal candidates for quantifying the effect of habitat filtering on the distribution and functional diversity of microscopic animals.

### Habitats and sampling design

As a study area, we selected the exposed seagrass meadow of Cala del Cuartel, in Santa Pola, south-eastern Spain (38° 12’ 34.04’’ N, 0° 30’ 19.12’’ W, WGS84 reference system), consisting of replicated patches at 4–7 m depth separated by bare sandy tongues. Marine mites prefer the *P. oceanica* patches and are rarely found in the sand (García-Gómez et al., submitted). So, in relation to the size and dispersal capabilities of the marine mites, each patch represents a discrete and independent replica of the same habitat within a larger area. The fact that all the patches are within the same bay limits the confounding effect of depth, temperature, salinity, or different exposition to currents.

Each patch consists of two compartments representing the two different habitats, the leaves and the matte (Figure 1A). The leaves are exposed to turbulence and affected by seasonal changes in length and growth of epiphytic algae and epifauna, which potentially represents the main source of food for the mites (Pugh & King, 1985a). In contrast, the matte is sheltered and offers a high and constant availability of detritus throughout the year.

**Figure 1.**
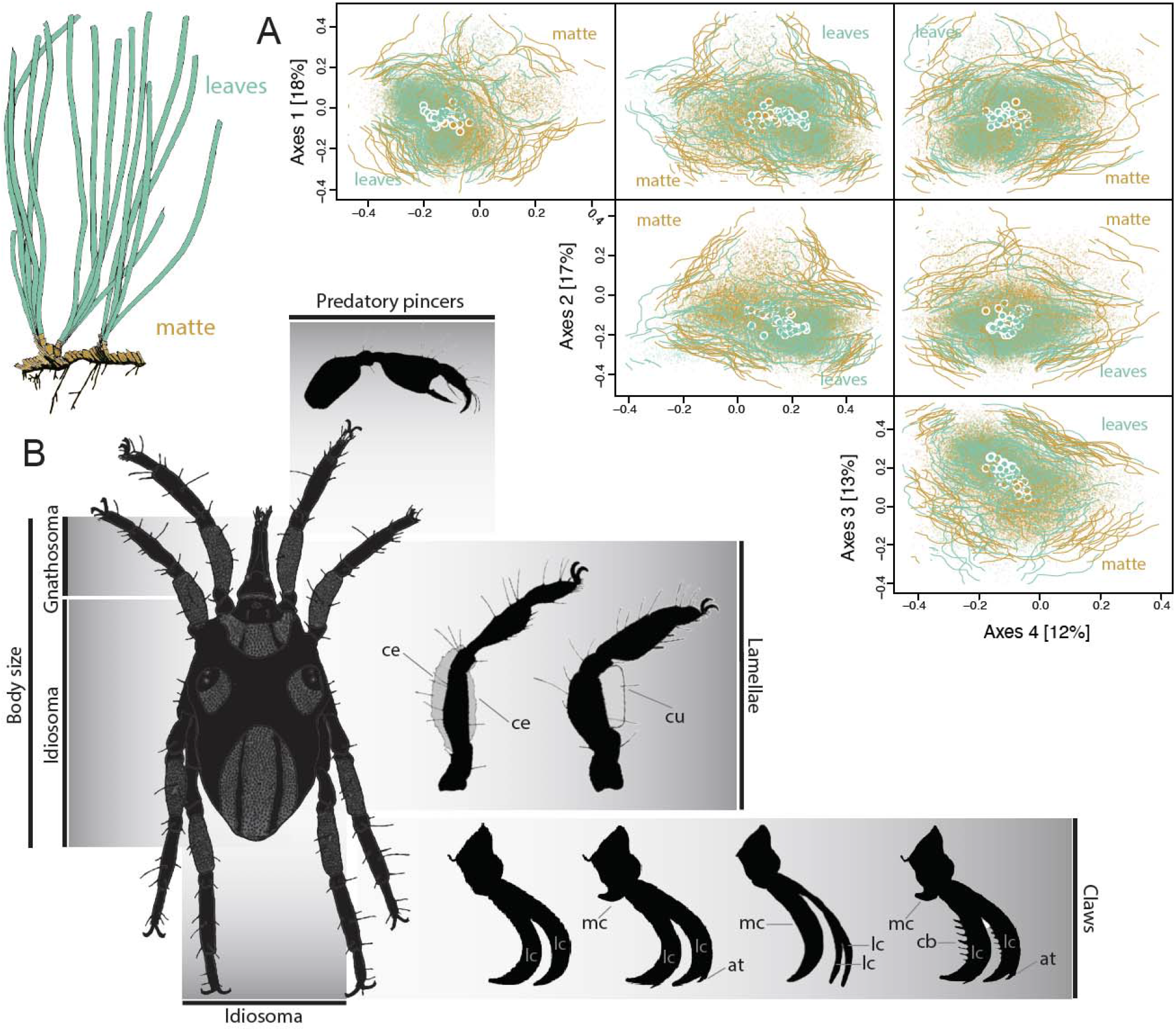
**A)** The 4-dimensional hypervolume of the mite communities in the *Posidonia oceanica* leaves (n=24) and matte (n=19). Large points with white borders represent the centroid of each hypervolume (note that due to the proximity of centroids, most points appear superimposed). The shape and boundaries of each hypervolume are defined by 1000 random points. All points are coloured according to the habitat. **B)** Summary of the morphological traits measured or estimated for each species and developmental stage. Further details on the interpretation of each trait are provided in Table 1 and the average values of traits across habitats in Figure S1. Abbreviations: *at* accessory tooth, *cb* comb, *ce* ceratogegumental lamellae, *cu* cuticular lamellae, *lc* lateral claw, *mc* median claw.

In each season between December 2015 and August 2016, scuba divers sampled these two habitats (leaves and matte) in six randomly selected patches of 400 cm^2^ of *Posidonia oceanica* (4 season x 6 patches x 2 habitats, totalling 48 samples). In each patch, leaves were collected first by cutting them at the ligulae level, while the surface of the underlying matte was scraped into a separate container.

**Table 1.** Morphological traits considered in the analyses, with hypotheses on their functional meaning

| Trait | Variable description | Functional meaning |
| --- | --- | --- |
| Total length | Measurement the tip of the gnathosoma to the tip of the idiosome in mm | Proxy of the total biovolume, trophic level and passive resistance of mites against water currents. |
| Idiosome length | Idiosome dorsal length | Proxy of the hard body length. |
| Idiosome width | Idiosome dorsal width | Proxy of the hard body width. |
| Gnathosoma (dorsal) length | Length of the gnathosoma which is not covered by the idiosome and exposed dorsally. | Proxy of the diet. The length of the gnathosoma is adapted to exploit different food resources (Bartsch 2006). |
| Idiosome length/width | Ratio between idiosome length and width | Proxy of body shape. Wider body shapes limit the colonization of habitat consisting of narrow spaces. Indeed, slender shaped mites are often found amongst fine sediments (Bartsch 2006). |
| Relative gnathosoma length | Ratio between gnathosoma dorsal length total body length | Proxy of the diet, as a measure of protruding gnathosoma relative to body size. |
| Accessory tooth | Categorical, reflecting the presence/absence of an accessory tooth on claws | In mites, especially those species linked to aquatic habitats, claws are essential to withstand physical stress, whether large (Pfingstl et al. 2020) or structural complex claws (Pugh & Fordy, 1987; Bartsch 2006). We here include four claw structures to account for different possible combinations |
| Combs | Degree of comb complexity, where 0 = absence, 1 = fine, 2 = regular, and 3 = large combs |  |
| Median claw type | Degree median claw development, where 0 = absence, 1 = small, and 2 = large median claw | that define claw complexity. The combination of these variables provides a proxy of the resistance of each individual to turbulence, as increasing claw complexity means a better grip to the substrate. |
| Number of legs with combs | Number of pairs of legs whose claws bear combs |  |
| Lamella | Categorical, reflecting the presence/absence of cerotegumental or cuticular lamella on legs | Lamella are present mostly in species that occur in sediments (Bartsch 2006). |
| Pincer | Categorical, reflecting the presence of a first pair of legs modified as a pincer | Specialised legs for feeding (Green & Macquitty 1987; Bartsch 2006). |

Meiofauna from each sample was extracted combining the magnesium chloride and the ‘bubble and blot’ decantation techniques to ensure the recovery of all species of marine mites (Higgins & Thiel, 1988; Sørensen & Pardos, 2008). The selected mesh size was 62 μm to collect both juveniles and adult forms. Each sample was bulk fixed using 7% formaldehyde in the field. All studied material has been deposited at the Laboratory of Meiofauna at the Universidad Complutense de Madrid.

In each habitat, we estimated a proxy for the availability of food. For each leaves sample, we estimated the average length of the leaves as the distance from the ligula to the apical end of all the complete leaves. Length of the leaves is known to correlate with the abundance of epiphytic organisms (Malbrouk, Hamza & Bradai, 2011). For each matte sample, we directly measured the percentage of organic carbon using the approach by Walkley & Black (1934).

### Species identification and morphological traits measurement

Mites were sorted using a MOTIC^®^ SMZ-168 stereoscope, whole-mounted in a modified Hoyer’s medium (Mitchell & Cook, 1952), and assigned to species and developmental stages by inspecting relevant morphological characters with a light microscope equipped with Nomarski optics and an Olympus DP70 camera. We used the keys by André (1946) and Green and MacQuitty (1987), as well as the available literature (Bartsch, 1991, 2000, 2001; Morselli, 1980).

For each species, we examined 13 morphological traits related to body size and shape, the ability to withstand the water currents, and trophic specialisation (Table 1). Body size and shape measures were taken on all 502 well-preserved specimens from our samples. The traits were estimated separately from adults and juveniles (larval or nymphal stages), as different life stages exhibit different ecological preferences and dispersal capabilities even within the same species (Bartsch, 2002; Somerfield & Jeal, 1995; 1996). The other traits, species-specific and not changing between individuals of different ages, were assigned at the species level.

### Functional space characterization

We expected the properties of the functional space to vary between the two different habitats, reflecting the habitat filtering effect in sorting the mite communities according to the presence of certain traits. Furthermore, we expected seasonal variations in the functional space in relation to the phenological changes of the *P. oceanica* meadow through the year. Therefore, we performed two sets of analyses: one, grouping all the samples from each habitat; and another, in which the samples were separated according to different surveys, each corresponding to a season.

We represented the functional space of mite communities in the two habitats and across seasons with geometrical *n*-dimensional hypervolumes (Blonder et al., 2014, 2018). Since some of the functional traits considered here are categorical, we applied a Gower dissimilarity measure to the complete trait matrix and extracted orthogonal morphological axes through principal coordinate analysis (Carvalho & Cardoso, 2020; Mammola & Cardoso, 2020). We delineated hypervolumes with the R package ‘*hypervolume’* (Blonder & Harris, 2018) using a gaussian kernel density estimate (Blonder et al., 2014, 2018), the first four principal coordinate axes (cumulatively 60% variance explained), a default bandwidth for each axis, and species abundances. A gaussian kernel density estimation was selected as it allows a probabilistic rather than a binary characterization of the functional space (Mammola & Cardoso, 2020). Five samples with one or no species were removed from the analyses. We analysed the properties of the hypervolumes with specific indices (Mammola & Cardoso, 2020) implemented in the R package ‘BAT’ (Cardoso, Rigal, & Carvalho, 2015; Cardoso, Mammola, Rigal, & Carvalho 2020). For each set of analyses, we expressed functional diversity with the *kernel*.*alpha* function as the total volume of the functional space. We verified if communities in matte and leaves and across seasons were subjected to different filtering processes by calculating the dispersion of the functional space with the *kernel*.*dispersion* function and the ‘divergence’ method (Mammola & Cardoso, 2020). The regularity of traits distributions within the total functional space was verified using the *kernel*.*evenness* function, which expresses evenness as the overlap between the input hypervolume and a theoretical hypervolume whose traits and abundances are evenly distributed within their possible range (Mammola & Cardoso, 2020).

We inspected whether certain assemblages of mite species act as indicators of the two habitats, and which species contribute most original traits to each habitat (i.e., functional outliers; Violle et al., 2017). In particular, we expect the distribution of the originality values to have a smaller variation in the leaves than in the matte, reflecting the stronger filtering effect exerted by this habitat compared to the matte. We calculated the functional originality of each species in each community with the function *kernel*.*originality*, weighting originality by species abundance (Mammola & Cardoso, 2020). We expressed originality as the average distance between each species to a sample of 10% stochastic points within the boundaries of the hypervolume. For each habitat, we expressed the total originality of a species as the average originality of the species across all communities in which it was present. Also, in this analysis, we considered the stages of the same species separately.

To define the degree to which a given species was characteristic to one habitat or the other, we further calculated the Δ Originality by subtracting to the value of originality of each species in the matte the value of originality of the same species in the leaves. When a species was absent in a habitat, we assigned its originality in this habitat to zero. We visualized Δ Originality values as histograms centred to the value of zero, where positive values indicate species that are more original in the matte than in the leaves, and negative values *vice versa*. We estimated and visualized the theoretical density of values with the R package ‘ggplot2’ (Wickham, 2016), by computing a kernel density estimate with a default bandwidth through the data.

To ease the interpretation of our findings, we finally calculated the probability of recovering a given trait within each habitat as the community weighted mean with the *cwm* function in ‘BAT’. For categorical traits, we calculated instead the probability of finding each state of the trait in each habitat using a function developed *ad hoc* for this study—see R code uploaded alongside this submission.

### Statistical analyses

We performed analysis of variance (ANOVA) to evaluate the significance of the differences observed in functional diversity, dispersion, and evenness between the matte and the leaves samples (Hypothesis 1), as well as amongst seasons (Hypothesis 3). When there was a significant effect of season, we performed a post hoc Tukey Honestly Significant Difference test to identify significant differences between pairs of seasons, using the R package ‘multcomp’ (Hothorn, Bretz, & Westfall, 2008). We verified whether the originality values of species in the leaves were significantly higher and lower than those in the matte using a null modelling approach (Hypothesis 2). We performed 99 permutations of the species between the two habitats, keeping fixed the original abundance values. For each run, we recalculated the hypervolumes and the originality values and estimated how many species in the leaves had higher originality than the species in the matte. As in Mammola et al. (2020), the null hypothesis of random sorting of species between the two habitats was rejected if the observed value was higher than the 97.5 percentile or lower than the 2.5 percentile of the 99 randomizations. For each permutation, we estimated the standard effect size and associated p-value.

## Results

We successfully reconstructed the hypervolumes for the 43 communities (that is, all those with more than one species). We observed a clear polarization of the trait space according to the two habitats (Figure 1). Properties of the functional space of the community in the two habitats were significantly different: the communities in the matte were functionally more diverse (ANOVA: F_(1,41)_ = 26.94, p < 0.001), more disperse (F_(1,41)_= 20.93, p < 0.001), and more even (F_(1,41)_ = 74.75, p < 0.001) than those in the leaves (Figure 2A, Table 2).

**Table 2.**
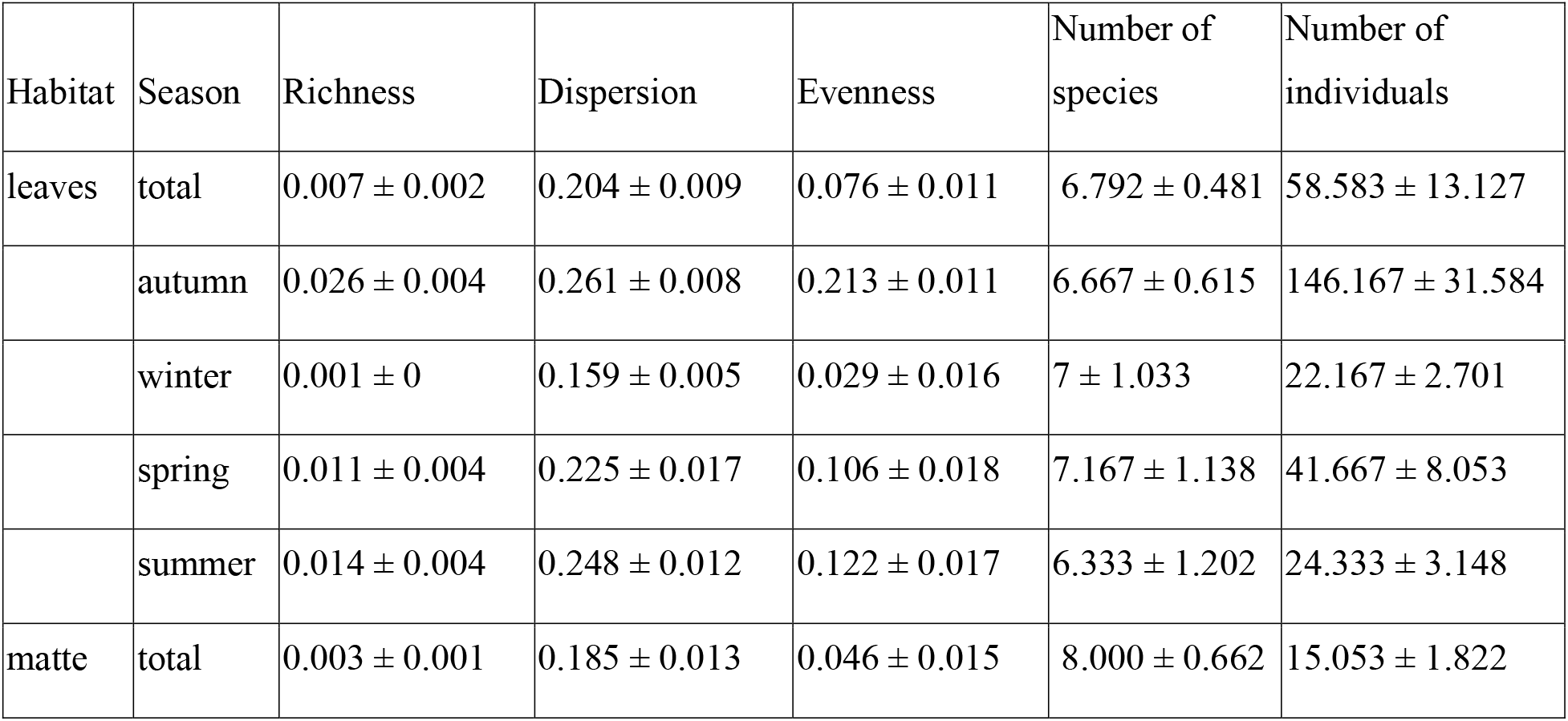
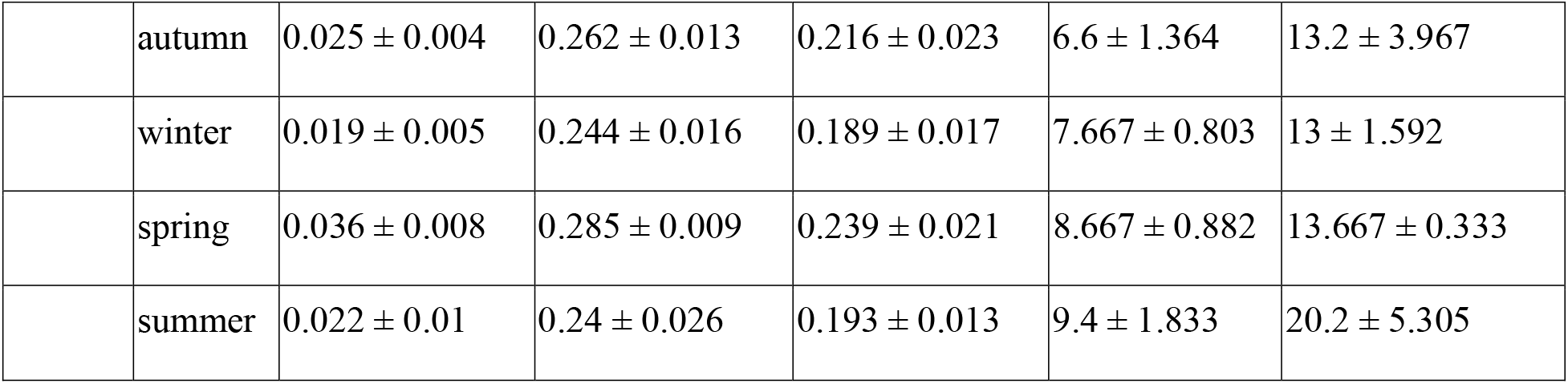
Summary of the average values (± standard error) of the number of species, number of individuals, and hypervolume metrics for the samples grouped by habitat (leaves and matte) and season.

**Figure 2.**
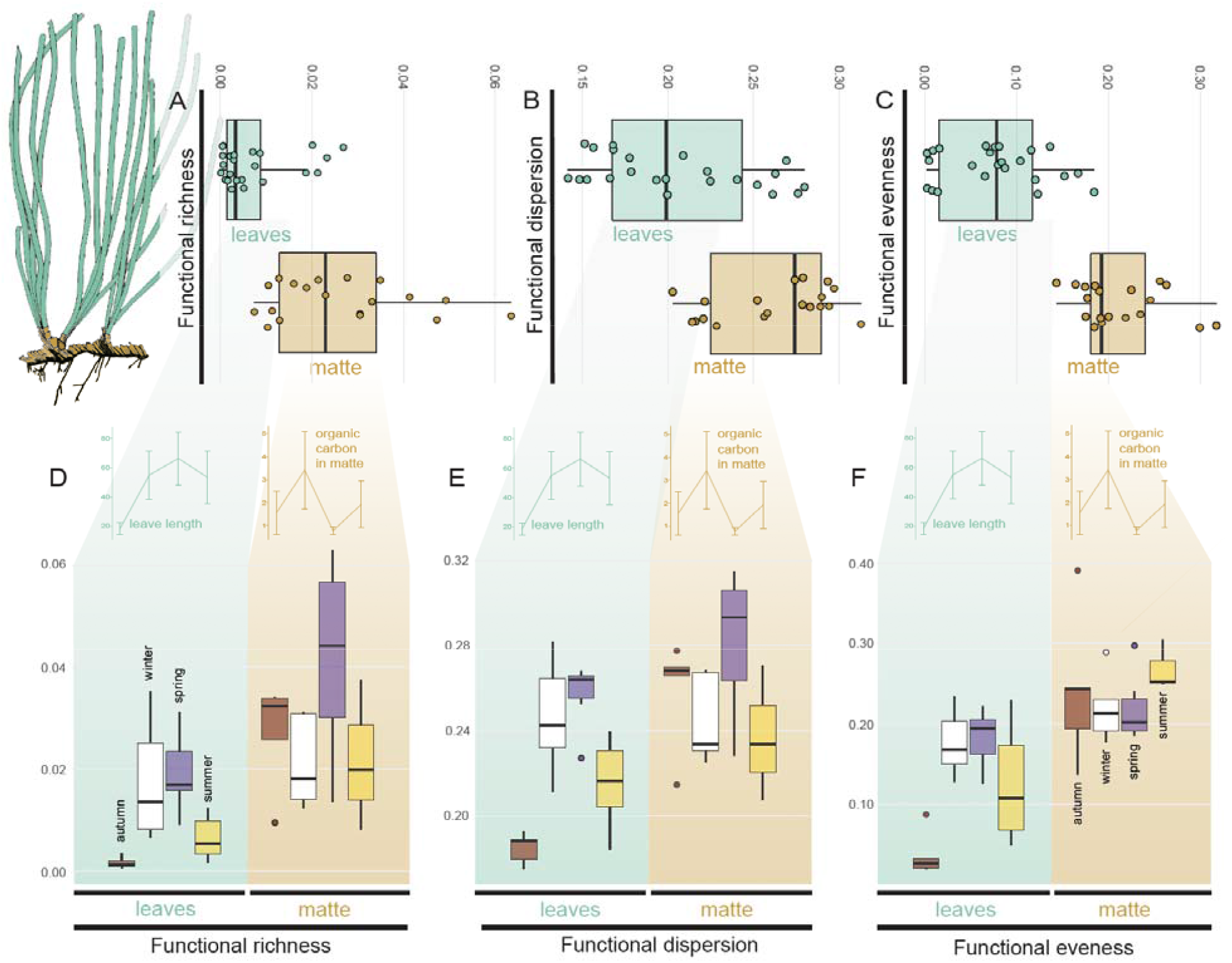
**A–C)** Overall differences in functional richness (A), dispersion (B) and evenness (C) between mite communities in leaves and matte. **D–F**) Differences in functional richness (D), dispersion (E) and evenness (F) across seasons. Inset graphs in d–f represent the variation in leaves mean length (in cm) for the leaves, and the organic matter content (in %) for the matte, thus reflecting the change in energy inputs due to the regeneration of leaves in the seagrass meadow across the four seasons.

Distribution of the total functional originality values was similar in both habitats (Figure 3A). According to the null modelling analysis, the number of species more original in the leaves than in the matter was not lower than what is expected from a random sorting of species across habitats (Standard effect size = –0.41, p-value = 0.06). Regarding the values of Δ Originality, we found a set of distinct species in the two habitats, allowing us to differentiate the leaves and matte communities according to the functional traits of few indicator species (Figure 3B).

**Figure 3.**
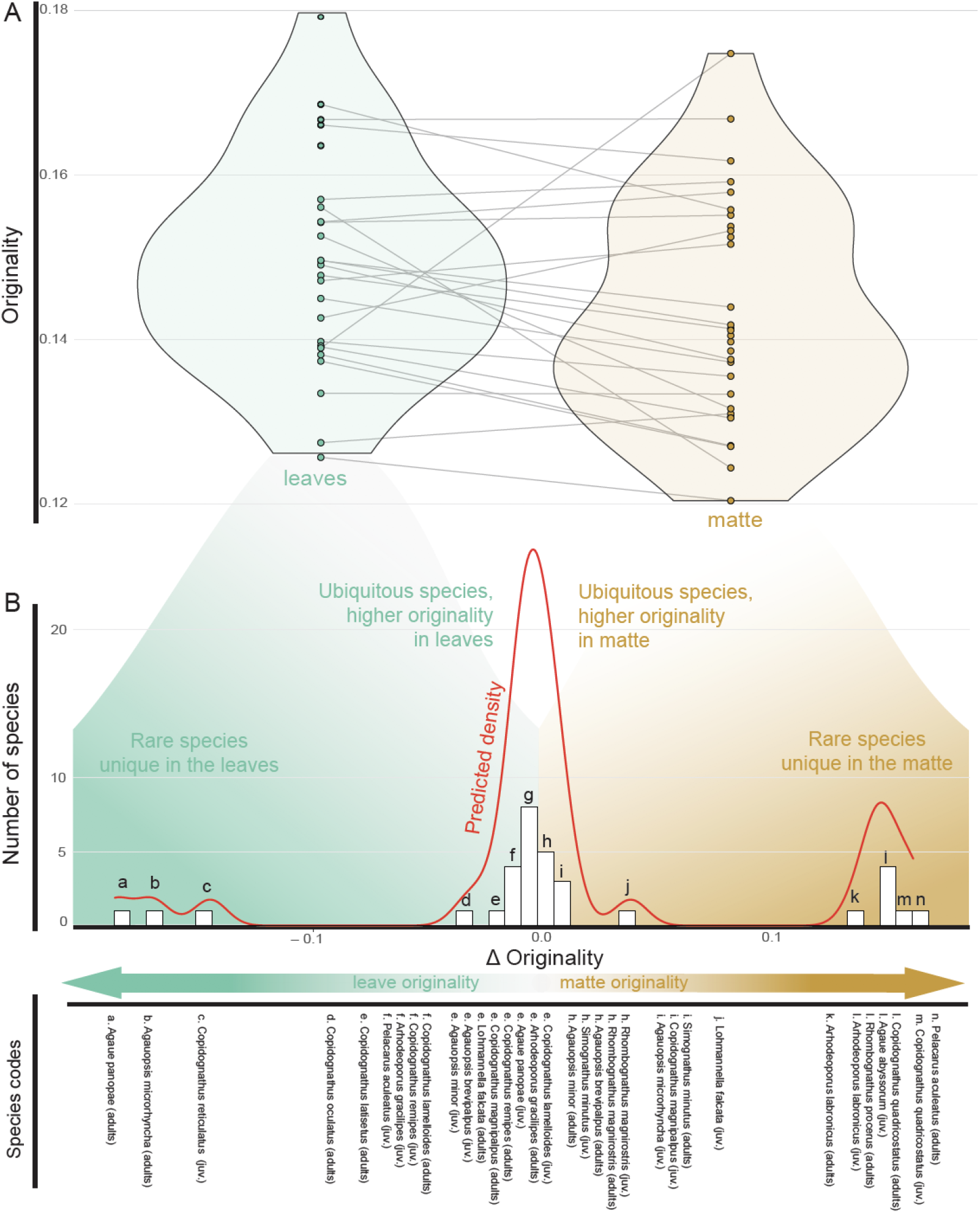
**A)** Violin plots showing the distribution of functional originality values of species in the leaves and the matte. Species present in both habitats are connected by grey lines. **B**) Histogram of Δ Originality values between species in the two habitats, calculated by subtracting the value of originality of each species in the leaves to the value of originality of each species in the matte. Orange smoothed lines show the predicted density of values according to a kernel density estimation. The letters above each bar correspond to the species listed at the rear of the figure.

There was a pronounced seasonal variability in the functional space of leave communities (Figure 2B), reflected in the differences in functional diversity (F_(3,20)_ = 5.146, p = 0.008), dispersion (F_(3,20)_ = 10.35, p < 0.001), and evenness (F_(3,20)_ = 7.593, p = 0.001) among seasons. In coincidence with the peaks of production of the meadow (Figure 2B, in-set graph), all three metrics were significantly higher in spring than in autumn and summer (Post-Hoc test: all p < 0.05). Functional dispersion and evenness were also significantly higher in winter than in autumn (Post-Hoc test: both p < 0.05). All other seasonal comparisons in the leaves were not significant (Post-Hoc test: all p > 0.05). In contrast, the seasonal pattern was not significant in the matte, neither for richness (ANOVA: F_(3,15)_ = 1.33, p = 0.303), nor for dispersion (F_(3,15)_ = 2.13, p = 0.139) nor evenness (F_(3,15)_ = 1.32, p = 0.306).

## Discussion

### Spatial patterns in functional diversity

Our analyses confirmed our first hypothesis that mite communities in matte habitat had a significantly higher functional richness, dispersion, and evenness than those in the leaves. Analytically, this means that, on average, the functional space in the leaves is significantly less voluminous (*i*.*e*. trait diversity is lower) and observations are less dispersed (*i*.*e*. species have traits that are more similar amongst them) and less even (*i*.*e*. the traits hypervolume is not homogenous indicating that certain combinations of traits are more common than others) than in the matte. Biologically, this suggests that the selective conditions in the leaves exert a stronger filtering effect upon the traits present in the colonizing species, whereby only a small subset from all the pool of traits present in the seagrass meadow allows mites to thrive in the leaves. This habitat filtering is reflected in the distribution of mites between habitats: even if the habitats are physically connected, communities in the leaves consist of a subset of the species present in the matte. The leaves are the habitat in which it is more likely to find individuals bearing specialised traits (Supplementary Material Figure S1). These traits are chiefly specialised claws (Figure S1d, S1e), which might aid in clinging to the leaf’s surface and thereby withstand turbulence (e.g. Pfingstl, Kerschbaumer, & Shimano, 2020; but see Pugh, King, & Fordy, 1987) and a larger body size (Figure S1g). In contrast, the assemblages in the matte consist of species bearing these traits, as well as species with more slender bodies (Figure S1i) and a longer and pointier gnathosoma (Figure S1j). Whereas the slender body presumably aids this species to crawl in the tighter habitat spaces in the matte, as observed in most interstitial microscopic species (Giere 2008), it is more difficult to interpret the functional meaning of the elongation of the gnathosoma. We here speculate that it might aid this species in feeding on detritus and deposits of organic matter accumulated in the tight spaces, but more in-depth studies would be needed to corroborate this assumption. A third group of species, presumably consisting of predators feeding on mites (Bartsch, 1989; J. Green & MacQuitty, 1987), are found occasionally in some of the samples, occurring stochastically both in the leaves and the matte as they wander around in the meadow searching for their prey.

This general pattern further emerges from the analysis of originality values, a metric that averages the distance between each observation to a sample of stochastic points within the boundaries of the hypervolume. It thereby measures how unique the position of individual observations is in the trait hyperspace, as the distances are expected to increase as the species’ combination of traits becomes unique (Mammola & Cardoso, 2020). Therefore, we expected more functionally original species in the matte, because species in the leaves need special adaptations presumably to cope with turbulence and feed on specialised food sources. The same adaptations are not required in the matte, where the presence of shelters and more diverse sources of food might relax the filtering effect on species and traits. This might result in a more functionally heterogeneous assemblage in which the probability of finding a given species is less dependent upon their traits. Our results, however, did not support this assumption given that originality values in the leaves did not differ significantly from those in the matte (Figure 3a). This might be the case because the species with the highest values of originality—such as *Pelacarus aculeatus, Agaue panopae, Agauopsis microrhyncha*, or *Agaue abyssorum*; Table S1—typically consisted of large rare species with uncommon traits that facilitate predation upon other microscopic animals, including mites (Bartsch, 1989; Green & MacQuitty, 1987). These species also occur in low abundances and their distribution is scattered across the meadow, being found stochastically in one habitat or the other. In fact, these species can be considered functional outliers (*sensu* Violle et al., 2017) in that they take extreme values of Δ Originality (Figure 3b), as they only occur in low numbers in either habitat, thus indicating that the filtering may act at another spatial or temporal scale on them. However, we acknowledge that further studies on the feeding biology of marine mites would be needed to fully understand the biological mechanisms behind the ecological patterns we documented.

### Temporal patterns in functional diversity

Our results partially corroborate our third hypothesis, as we found significant temporal variations in the functional diversity of mite communities in the leaves likely following the annual cycle of the *Posidonia oceanica*. As above, these changes permeate all metrics, which were significantly higher in spring than in autumn and summer, in coincidence with the spring peaks of production in the meadow. Functional dispersion and evenness were also significantly higher in winter than in autumn.

The end of the summer is characterized in the Mediterranean by an increase of the rainfall and primary production, which favours a rapid growth of *P. oceanica* in winter reaching a peak in the biomass in the seagrass meadow in spring (Champenois & Borges, 2014). A large number of epiphytes colonize the leaves, which get densely populated by diverse epiphytic communities (Mabrouk, Hamza, Brahim, & Bradai, 2011; Piazzi, Balata, & Ceccherelli, 2016), as they enlarge. Food resources are hence more abundant and diverse in the leaves at their peak of production in spring, which positively feedbacks the mite populations. Furthermore, the basal parts of long leaves are less exposed to hydrodynamics, as leaves themselves provide shelter from the current towards the bottom (Folkard, 2005). These two factors, increase of food and higher shelter, presumably result in a milder ecological filter, enhancing the possibility for different mites to exploit this habitat and reproduce therein. Indeed, juveniles, which have not developed yet all their adult traits to withstand currents (e.g. smaller body or legs with fewer segments, yet provided with claws as in adults), become dominant in the long leaves exclusively in spring (García-Gómez et al., submitted).

In contrast, the matte does not experience similar pronounced phenological changes and we can speculate that this is the reason for which no significant changes were observed in the functional diversity of mite communities in the matte.

### Conclusion

Being the first study using hypervolumes to define functional properties of meiofauna communities, our study highlights a potential role of the environment in affecting the distribution of microscopic animals between connected habitats by filtering them according to the presence of certain traits. Remarkably, this filtering effect was relatively weak, as most species were found in both habitats and the filtering was mostly reflected by their relative abundances. Therefore, one may argue that our results of filtering effects between connected habitats might not be applied to all microscopic animals more widely and that mites in seagrass meadows might represent only a specific case. Similar filtering effects might be even more subtle and difficult to isolate in other microscopic animal groups (rotifers, tardigrades, and soft-bodied groups) for which the functional interpretation of morphological traits is often obscure and trait measurements subjected to strong artefacts due to post-mortem contraction, fixation, and other bias (Higgins & Thiel, 1988). Furthermore, most microscopic animals have a high probability to be passively dispersed to suboptimal habitats (Armonies, 1988; Hagerman & Rieger, 1981; Hauspie & Polk, 1973), increasing the uncertainty associated with habitat characterization at a small scale relevant for their biology, thus overestimating their potential Grinnellian niche.

Therefore, it is not surprising that in such studies the distribution of microscopic animals might appear either uniform or random, simply as a consequence of the high uncertainty associated with measurements and morphological interpretation at the small spatial scales. In other words, microscopic size may generate uncertainty in a macroscopic observer, on both the definition of traits and the definition of niche even if *the environment did select*. Exploring the distribution of small animals through the lens of functional ecology, targeting traits with clear functional meaning related to habitat occupation, is crucial to overcome some of these biases (Violle, Reich, Pacala, Enquist, & Kattge, 2014). Our study therefore emphasises the need of moving from a merely taxonomical toward a functional view of ecological studies of microscopic organisms (Green, Bohannan, & Whitaker, 2008).

## Acknowledgements

The authors want to thank Dr Alfonso Ramos-Esplá (CIMAR) for the logistical assistance during the sampling campaigns in Santa Pola. We also thank Dr Sergio Pérez and Nuria Rico for the help with the samplings. We gratefully acknowledge Dr Ilse Bartsch for her invaluable help with the identification of halacarid species. We also thank Dr Dolores Trigo and María Isla for the facilities and their useful assistance in the laboratory of soil chemistry of the Zoology Department (UCM). SM was supported by the CAWEB project “Testing macroecological theory using simplified systems”, funded by the European Commission through Horizon 2020 Marie Skłodowska-Curie Actions (MSCA) individual fellowships (Grant no. 882221). GGG was supported by an Erasmus+ mobility fellowship, OLS ID 641798. NS was funded by the Research Talent Attraction Program (Regional Government of Madrid and University Complutense of Madrid) (2019-T2/AMB-13328).

## Supplementary materials online

**Table S1**. Summary traits, number of examined individuals, and originality values per species and life stage. The definition of each trait is included in Table 1.

## Supplementary material Figure S1

**Figure S1.**
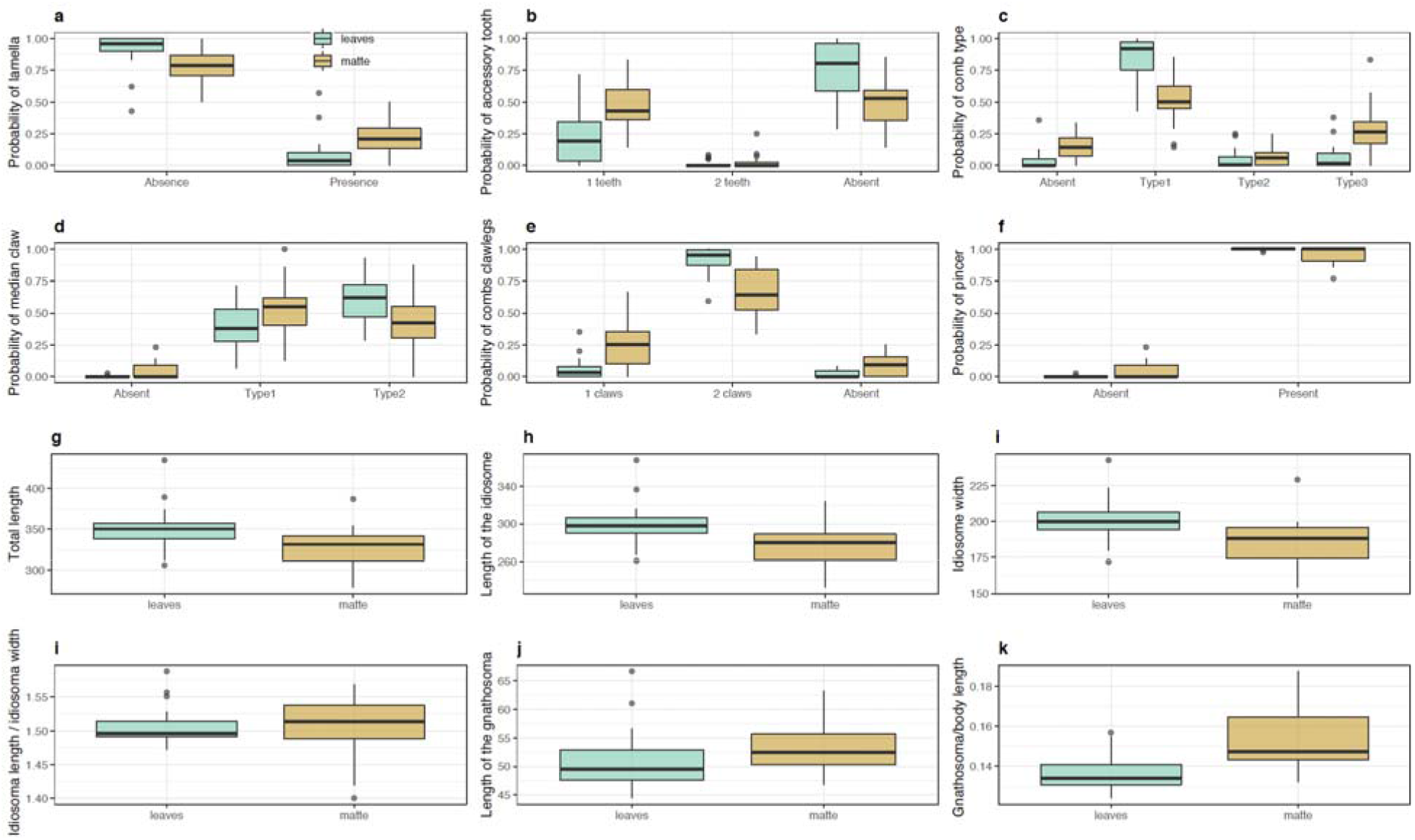
Probability of finding each state of discrete traits **(a–f)** and community weighted mean of continuous traits **(g–k)** for mite communities in the leaves and matte.

## REFERENCES

Adams, D. (1984). Life, the Universe and Everything: Hitchhiker’s Guide to the Galaxy Book 3 (Vol. 3). Tor UK.

André, M. (1946). Halacariens marins. Faune de France, 46, 1–152.

Armonies, W. (1988). Active emergence of meiofauna from intertidal sediment. Marine Ecology Progress Series, 43, 151–159. doi: 10.3354/meps043151

Baas-Becking, L. G. M. (1934). Geobiologie; of inleiding tot de milieukunde. WP Van Stockum & Zoon NV.

Bartels, P. J., Kaczmarek, L., Rozkowska, M., & Nelson, D. (2020). Interactive map of marine tardigrades of the world. https://paul-bartels.shinyapps.io/marine-tardigrades/.

Bartsch, I. (1989). Marine mites (Halacaroidea: Acari): a geographical and ecological survey. Hydrobiologia, 178(1), 21–42.

Bartsch, I. (1991). Taxonomic notes on halacarids (Acari) from the Skagerrak area. Helgoländer Meeresuntersuchungen, 45(1–2), 97–106.

Bartsch, I. (2000). A new species of *Isobactrus* from the Black Sea (Acari, Halacaridae, Rhombognathinae). Cahiers de Biologie Marine, 41(4), 407–412.

Bartsch, I. (2001). Black Sea Copidognathinae (Arachnida, Acari, Halacaridae): a review. Zoosystematics and Evolution, 77(2), 247–275.

Bartsch, I. (2002). Geographical and ecological distribution of marine halacarid genera and species (Acari: Halacaridae). Experimental & Applied Acarology, 34(1–2), 37–58.

Bass, D., & Boenigk, J. (2011). Everything is everywhere: a twenty-first century de-/reconstruction with respect to protists. Biogeography of Microscopic Organisms: Is Everything Small Everywhere, 88–110.

Blonder, B, & Harris, D. (2018). High dimensional geometry and set operations using kernel density estimation, support vector machines, and convex hulls. R Package, 2, 11.

Blonder, B., Lamanna, C., Violle, C., & Enquist, B. J. (2014). The *n*-dimensional hypervolume. Global Ecology and Biogeography, 23(5), 595–609. doi: 10.1111/geb.12146

Blonder, B., Morrow, C. B., Maitner, B., Harris, D. J., Lamanna, C., Violle, C., Enquist, B.J., Kerkhoff, A. J. (2018). New approaches for delineating *n* Ldimensional hypervolumes. Methods in Ecology and Evolution, 9(2), 305–319. doi: 10.1111/2041-210X.12865

Borgonie, G., García-Moyano, A., Litthauer, D., Bert, W., Bester, A., van Heerden, E., Erasmus, M., Onstott, T. (2011). Nematoda from the terrestrial deep subsurface of South Africa. Nature, 474(7349), 79–82.

Cardoso, P., Rigal, F., & Carvalho, J. C. (2015). BAT–Biodiversity Assessment Tools, an R package for the measurement and estimation of alpha and beta taxon, phylogenetic and functional diversity. Methods in Ecology and Evolution, 6(2), 232–236.

Carvalho, J. C., & Cardoso, P. (2020). Decomposing the causes for niche differentiation between species using hypervolumes. Frontiers in Ecology and Evolution, 8, 243.

Champenois, W., & Borges, A. (2014). Seasonal and inter-annual variations of community metabolism rates of a Posidonia oceanica seagrass meadow based on continuous oxygen measurements with optodes.

Drew, E. A. (1978). Factors affecting photosynthesis and its seasonal variation in the seagrasses *Cymodocea nodosa* (Ucria) Aschers, and *Posidonia oceanica* (L.) Delile in the Mediterranean. Journal of Experimental Marine Biology and Ecology, 31(2), 173–194.

Folkard, A. M. (2005). Hydrodynamics of model *Posidonia oceanica* patches in shallow water. Limnology and Oceanography, 50(5), 1592–1600.

Fontaneto, D., Westberg, M., & Hortal, J. (2011). Evidence of weak habitat specialisation in microscopic animals. PLoS One, 6(8), e23969. doi: 10.1371/journal.pone.0023969

Fonseca, G., & Netto, S. A. (2015). Macroecological patterns of estuarine nematodes. Estuaries and Coasts, 38(2), 612–619. doi: 10.1007/s12237-014-9844-z

Fontaneto, D. (2019). Long-distance passive dispersal in microscopic aquatic animals. Movement Ecology, 7(1), 10. doi: 10.1186/s40462-019-0155-7

Fontaneto, D., Barraclough, T. G., Chen, K., Ricci, C., & Herniou, E. A. (2008). Molecular evidence for broad-scale distributions in bdelloid rotifers: everything is not everywhere but most things are very widespread. Molecular Ecology, 17(13), 3136–3146. doi: 10.1111/j.1365-294X.2008.03806.x

Fontaneto, D., & Hortal, J. (2013). At least some protist species are not ubiquitous. Molecular Ecology, 22(20), 5053–5055. doi: 10.1111/mec.12507

García-Gómez, G., García-Herrero, Á., Sánchez, N., Pardos, F., Izquierdo-Muñoz, A., Fontaneto, D., & Martínez, A. (submitted). Ecological preferences shift the distribution of marine mites in posidonia oceanica seagrass.

Giere, O. (2008). Meiobenthology: the microscopic motile fauna of aquatic sediments. Springer Science & Business Media.

Green, J. L., Bohannan, B. J., & Whitaker, R. J. (2008). Microbial biogeography: from taxonomy to traits. Science, 320(5879), 1039–1043.

Green, J., & MacQuitty, M. (1987). Halacarid Mites (Arachnida: Acari) keys and notes for the identification of the species, Synopses of the British Fauna, ed: Kermak, DM and Barnes, RSK No: 36. The Linnean Society, London, 178p.

Hagerman, G. M., & Rieger, R. M. (1981). Dispersal of benthic meiofauna by wave and current action in Bogue Sound, North Carolina, USA. Marine Ecology, 2(3), 245–270.

Hauspie, R., & Polk, P. (1973). Swimming behaviour patterns in certain benthic harpacticoids (Copepoda). Crustaceana, 25(1), 95–103.

Higgins, R. P., & Thiel, H. (1988). Introduction to the study of meiofauna. Washington, D.C: Smithsonian Institution Press.

Hoschitz, M., & Kaufmann, R. (2004). Soil nematode communities of Alpine summits–site differentiation and microclimatic influences. Pedobiologia, 48(4), 313–320. doi: 10.1016/j.pedobi.2004.03.004

Hothorn, T., Bretz, F., & Westfall, P. (2008). Simultaneous inference in general parametric models. Biometrical Journal: Journal of Mathematical Methods in Biosciences, 50(3), 346–363.

Kaczmarek, L., Michalczyk, Ł., & McInnes, S. J. (2015). Annotated zoogeography of non-marine Tardigrada. Part II: South America. Zootaxa, 3923(1), 1–107.

Levin, S. A. (1992). The problem of pattern and scale in ecology: the Robert H. MacArthur Award Lecture. Ecology, 73(6), 1943–1967. doi: 10.2307/1941447

Mabrouk, L., Hamza, A., Brahim, M. B., & Bradai, M. (2011). Temporal and depth distribution of microepiphytes on *Posidonia oceanica* (L.) Delile leaves in a meadow off Tunisia. Marine Ecology, 32(2), 148–161.

Mammola, S., Arnedo, M. A., Fišer, C., Cardoso, P., Dejanaz, A. J., & Isaia, M. (2020). Environmental filtering and convergent evolution determine the ecological specialization of subterranean spiders. Functional Ecology, 34(5), 1064–1077.

Mammola, S., Amorim, I. R., Bichuette, M. E., Borges, P. A. V., Cheeptham, N., Cooper, S. J. B., Culver, D.C., Deharveng, L., Eme, D., Ferreira, R.L., Fišer, C., Fišer, Ž., Fong, D.W., Griebler, C., Jeffery, W.R., Jugovic, J., Kowalko, J.E., Lilley, T.M., Malard, F., Manenti, R., Martínez, A., Meierhofer, M.B., Niemiller, M.L., Northup, D.E., Pellegrini, T.G., Pipan, T., Protas, M., Reboleira, A.S.P.S., Venarsky, M.P., Wynne, J.J., Zagmajster, M., Cardoso, P. (2020). Fundamental research questions in subterranean biology. Biological Reviews, brv.12642. doi: 10.1111/brv.12642

Mabrouk, L., Hamza, A., Brahim, M.B., & Bradai, M.N. (2011). Temporal and depth distribution of microepiphytes on *Posidonia oceanica* (L.) Delile leaves in a meadow off Tunisia. Marine Ecology, 32: 148–16. doi.org/10.1111/j.1439-0485.2011.00432.x

Mammola, S., & Cardoso, P. (2020). Functional diversity metrics using kernel density *n* LJdimensional hypervolumes. Methods in Ecology and Evolution, 2041-210X.13424. doi: 10.1111/2041-210X.13424

Martínez, A., Di Domenico, M., Leasi, F., Curini-Galletti, M., Todaro, M.A., Zotto, M.D., Gobert, S., Artois, T., Norenburg, J., Jörger, K.M., Núñez, J., Fontaneto, D., Worsaae, K. (2019). Patterns of diversity and endemism of soft-bodied meiofauna in an oceanic island, Lanzarote, Canary Islands. Marine Biodiversity, 49(5), 2033–2055. doi: 10.1007/s12526-019-01007-0

Mitchell, R. D., & Cook, D. R. (1952). The preservation and mounting of water-mites. Turtox News, 30(9), 1–4.

Molenaar, H., Barthélémy, D., De Reffye, P., Meinesz, A., & Mialet, I. (2000). Modelling architecture and growth patterns of *Posidonia oceanica*. Aquatic Botany, 66(2), 85–99.

Morselli, I. (1980). Su tre acari prostigmati di acque salmastre dell’alto Adriatico. Atti Della Societa Toscana Di Scienza Naturali Memorie, Serie B, 87, 181–195.

Ott, J., Rieger, G., Rieger, R., & Enderes, F. (1982). *Astomonema jenneri*, a new mouthless nematode and the evolution of the association between prokaryotes and interstitial worms. Marine Ecology, 3(4), 313–333.

Pepato, A. R., Vidigal, T. H., & Klimov, P. B. (2018). Molecular phylogeny of marine mites (Acariformes: Halacaridae), the oldest radiation of extant secondarily marine animals. Molecular Phylogenetics and Evolution, 129, 182–188.

Pfingstl, T., Kerschbaumer, M., & Shimano, S. (2020). Get a grip—evolution of claw shape in relation to microhabitat use in intertidal arthropods (Acari, Oribatida). PeerJ, 8, e8488.

Piazzi, L., Balata, D., & Ceccherelli, G. (2016). Epiphyte assemblages of the Mediterranean seagrass Posidonia oceanica: an overview. Marine Ecology, 37(1), 3–41.

Potter, K. A., Arthur Woods, H., & Pincebourde, S. (2013). Microclimatic challenges in global change biology. Global Change Biology, 19(10), 2932–2939.

Pugh, P., & King, P. (1985a). Feeding in intertidal Acari. Journal of Experimental Marine Biology and Ecology, 94(1–3), 269–280.

Pugh, P., & King, P. (1985b). Vertical distribution and substrate association of the British Halacaridae. Journal of Natural History, 19(5), 961–968.

Pugh, P., King, P., & Fordy, M. (1987). Possible significance of the claw structure in the Rhombognathinae (Halacaridae, Prostigmata, Acari). Acarologia, 28(2), 171–175.

Soberón, J., & Nakamura, M. (2009). Niches and distributional areas: Concepts, methods, and assumptions. Proceedings of the National Academy of Sciences, 106(Supplement 2), 19644–19650. doi: 10.1073/pnas.0901637106

Somerfield, P. J., & Jeal, F. (1996). Vertical distribution and substratum association of Halacaridae (Acari: Prostigmata) on sheltered and exposed Irish shores. Oceanographic Literature Review, 1(43), 62.

Somerfield, P., & Jeal, F. (1995). Vertical distribution and substratum association of Halacaridae (Acari: Prostigmata) on sheltered and exposed Irish shores. Journal of Natural History, 29(4), 909–917.

Sørensen, M. V., & Pardos, F. (2008). Kinorhynch systematics and biology – an introduction to the study of kinorhynchs, inclusive identification keys to the genera. Meiofauna Marina, 16, 21–73.

Violle, C., Reich, P. B., Pacala, S. W., Enquist, B. J., & Kattge, J. (2014). The emergence and promise of functional biogeography. Proceedings of the National Academy of Sciences, 111(38), 13690–13696.

Violle, C., Thuiller, W., Mouquet, N., Munoz, F., Kraft, N. J., Cadotte, M. W., Livingstone, S.W., Mouillot, D. (2017). Functional rarity: the ecology of outliers. Trends in Ecology & Evolution, 32(5), 356–367.

Walkley, A., & Black, I. A. (1934). An examination of the Degtjareff method for determining soil organic matter, and a proposed modification of the chromic acid titration method. Soil Science, 37(1), 29–38.

Wickham, H. (2016). ggplot2: elegant graphics for data analysis. springer.

Worsaae, K., Gonzalez, B. C., Kerbl, A., Nielsen, S. H., Jørgensen, J. T., Armenteros, M., Iliffe, T.M., Martínez, A. (2019). Diversity and evolution of the stygobitic *Speleonerilla* nom. nov. (Nerillidae, Annelida) with description of three new species from anchialine caves in the Caribbean and Lanzarote. Marine Biodiversity, 49(5), 2167–2192. doi: 10.1007/s12526-018-0906-5

Zeppilli, D., Leduc, D., Fontanier, C., Fontaneto, D., Fuchs, S., Gooday, A. J., Gooday, Goinea A., Ingels, J., Ivanenko, V.N., Kristensen, R.M., Neves, R.C., Sánchez, N., Sandulli, R., Sarrazin, J., Sørensen, M.V., Tasiemski, A., Vanreusel, A., Autret, M., Bourdonnay, L., Claireaux, M., Coquillé, V., De Wever, L., Rachel, D., Marchant, J., Toomey, L. & Fernandes, D. (2018). Characteristics of meiofauna in extreme marine ecosystems: a review. Marine Biodiversity, 48(1), 35–71. doi: 10.1007/s12526-017-0815-z

Zupo, V., Buia, M., & Mazzella, L. (1997). A production model for *Posidonia oceanica* based on temperature. Estuarine, Coastal and Shelf Science, 44(4), 483–492.

